# FENNEC: photon-level deep learning for classifying bursts in diffusion-based single-molecule FRET

**DOI:** 10.64898/2026.07.14.737247

**Authors:** Bob Schiffrin, Joel A. Crossley

## Abstract

Single-molecule Förster resonance energy transfer (smFRET) reports on biomolecular conformational dynamics by measuring distance changes between donor and acceptor fluorophores. In diffusion-based smFRET, however, the detection of genuine conformational exchange is routinely confounded by photophysical artefacts, notably acceptor photobleaching and blinking, which produce similar burst-level signatures. Many existing methods for resolving conformational dynamics and dye photophysics rely on fitting kinetic models with a fixed number of states, which is typically unknown. Here we present FENNEC (Fluorescence Event Neural Network for Evaluating and Classifying bursts), a dilated convolutional neural network for diffusion-based smFRET data that simultaneously detects conformational dynamics, acceptor photobleaching, and acceptor blinking within individual bursts, directly from raw photon arrival times. FENNEC is trained entirely on simulated data, and requires no experimental data with assigned labels for training. Crucially, the dynamics classification is independent of the number of underlying states, and therefore provides an analysis and filtering method that complements established methods that extract the number of states and their kinetics. FENNEC can identify a high-confidence subset of static and dynamic bursts, while ambiguous bursts can be excluded or set aside for further analysis. Applied to a dynamic DNA hairpin, FENNEC recovers the expected population distributions. Together, these results provide proof of principle that a classifier trained on simulated photon-level data can identify conformational dynamics and photophysical artefacts in experimental smFRET data, and we invite further evaluation on a range of instruments and systems. FENNEC is freely available at https://github.com/jacrossley/FENNEC.

## Introduction

Single-molecule Förster resonance energy transfer (smFRET) measures distances between different regions of biomolecules at the nanometre scale (∼2-10 nm), providing direct information on conformational states and dynamics in solution (Ha et al., 2024; Lerner et al., 2021; Lerner, Cordes, et al., 2018; Mazal & Haran, 2019; Nettels et al., 2024). In a typical experiment, donor and acceptor fluorophores are attached to a biomolecule of interest at defined sites, and the efficiency (E) of energy transfer between the pair is monitored within individual molecules over time. This can provide information on structural heterogeneity and interconversion kinetics, and has been applied to processes including protein folding (Chung, 2018; Schuler & Eaton, 2008), ligand binding (Husada et al., 2018; Schiffrin et al., 2024), and allosteric regulation (Crossley et al., 2024; Liauw et al., 2022; Mazal & Haran, 2019). Diffusion-based smFRET, in which freely diffusing molecules traverse a confocal excitation volume, allows the measurement of thousands of individual molecules per hour to give reliable population statistics, while avoiding potential surface immobilisation artefacts. When combined with alternating laser excitation (ALEX) (Kapanidis et al., 2004) or pulsed interleaved excitation (PIE) (Müller et al., 2005), diffusion-based smFRET provides per-burst stoichiometry (S) information, allowing molecules containing only a single fluorophore, or in which only one fluorophore is currently fluorescing, to be filtered out. Individual fluorescence bursts typically last ∼1 ms, during which only a relatively small number of photons (tens to hundreds) are detected. A key challenge in analysing this burst data is separating bursts containing a single FRET state (e.g. molecules residing in one conformational state) from those which sample multiple FRET states (e.g. molecules undergoing conformational exchange), a distinction made difficult by the noisy, stochastic nature of single-molecule data. This problem is compounded by photophysical artefacts that produce fluorescence signatures closely resembling genuine FRET dynamics: acceptor photobleaching causes an abrupt, irreversible loss of FRET signal that mimics a transition to a low-FRET state, while transient acceptor blinking produces intermittent drops in acceptor emission, and hence apparent low FRET, on timescales that overlap with conformational exchange itself.

Conformational dynamics within individual bursts are currently detected with a variety of analytical approaches. Burst variance analysis (BVA) compares the observed variance in FRET efficiency to that expected from shot noise alone (Torella et al., 2011). The two-channel kernel density estimator (FRET-2CDE) uses the relative timing of donor and acceptor photons to identify bursts in which FRET efficiency changes over time (Tomov et al., 2012). In pulsed excitation schemes, E-τ analysis identifies dynamics as deviations from the fixed relationship between mean FRET efficiency and the donor fluorescence lifetime expected for a single FRET state (Maus et al., 2001; Widengren et al., 2006). These methods can detect changes in FRET efficiency during a burst, but they struggle to distinguish genuine conformational dynamics from photophysical artefacts of the fluorophores, which can lead to misleading conclusions (Bagshaw & Cherny, 2006; Roy et al., 2008). More model-dependent approaches such as multi-parameter photon-by-photon hidden Markov modelling (mpH^2^MM) enable explicit kinetic modelling of state interconversion within bursts and can help filter out photophysical artefacts (Harris et al., 2022; Kache & Hendrix, 2025; Pirchi et al., 2016), but require the number of underlying states to be specified in advance for each model fit. Model selection can be aided by integrated completed likelihood (Harris et al., 2022) or modified Bayesian information criterion (Lerner, Ingargiola, et al., 2018). However, these criteria can often disagree, and determining the correct number of states typically requires prior knowledge that is difficult to obtain when the system under study is poorly characterised.

Machine learning is increasingly used in the physical and life sciences to analyse complex data (Vinuesa et al., 2026). Within single-molecule biophysics specifically, machine learning approaches have been applied successfully to step detection in camera-based surface-immobilised fluorescence traces (Thomsen et al., 2020; Wanninger et al., 2023), single-molecule fluorescence localisation microscopy (Zhang et al., 2023), and the classification of molecular trajectories from fluorescence tracking experiments (Pambos et al., 2026). WaveNet-style neural networks based on dilated convolutions, originally developed for the generative modelling of raw audio waveforms (van den Oord et al., 2016), are particularly well-suited to long, sparse, sequential data with structure on multiple timescales. Exponentially increasing dilations provide a receptive field that grows with depth, allowing the network to capture both fine-grained local patterns and long-range dependencies across thousands of input positions. This is computationally efficient as it avoids the quadratic scaling with input length associated with attention-based models such as transformers (Bai et al., 2018). Photon streams within bursts are inherently sequential, with stochastic inter-photon arrival times, and encode underlying photophysical and conformational processes spanning several orders of magnitude (μs-ms). This led us to hypothesise that WaveNet-style dilated convolutional neural networks (CNNs), which are well suited to long-range sequential dependencies, might offer a natural fit for classifying individual smFRET bursts.

Here we present FENNEC (Fluorescence Event Neural Network for Evaluating and Classifying bursts), a dilated convolutional deep learning model that simultaneously identifies conformational dynamics, acceptor photobleaching, and acceptor blinking within individual smFRET bursts directly from raw photon arrival times. FENNEC is trained entirely on simulated data calibrated to the user’s instrument and sample, avoiding the need for experimentally labelled training sets. Crucially, the dynamic/static classification is agnostic to the number of underlying conformational states, providing a filter to remove bursts that contain photophysical artefacts and complementing modelling approaches such as mpH^2^MM (Harris et al., 2022; Pirchi et al., 2016) without requiring the number of states to be specified in advance. We apply FENNEC to a canonical two-state dynamic smFRET system, a DNA hairpin (Harris et al., 2022; Tomov et al., 2012; Tsukanov et al., 2013), as a proof of principle. FENNEC recovers physically interpretable population distributions, isolates dynamic bursts for potential downstream kinetic analysis, and identifies acceptor blinking/bleaching in experiments that would otherwise confound dynamics detection. Together, these results establish FENNEC as a burst classifier that identifies dynamic bursts, and separates them from bursts affected by acceptor bleaching or blinking, directly from raw photon arrival times.

## Results

### FENNEC classifies dynamics, blinking and bleaching using instrument-calibrated simulated training data

In diffusion-based smFRET experiments, freely diffusing molecules produce brief fluorescence bursts as they traverse a confocal volume (**Figure 1a**). In ALEX or PIE measurements, each photon within a burst is recorded with information about the excitation source and the emission detection channel, giving four photon streams: donor-excitation with donor emission (D_ex_D_em_), donor-excitation with acceptor emission (D_ex_A_em_), acceptor-excitation with acceptor emission (A_ex_A_em_), and acceptor-excitation with donor emission (A_ex_D_em_). The D_ex_A_em_ stream monitors acceptor emission following donor excitation and is therefore the main FRET reporter. Here, only three of these are used: D_ex_D_em_, D_ex_A_em_, and A_ex_A_em_, as A_ex_D_em_ is not informative for the classification tasks considered here (**Figure 1b**). FENNEC operates at the single-photon level, where the raw photon streams in each channel carry information about the underlying FRET states (**Figure 1c**). A central challenge is that genuine conformational dynamics, acceptor photobleaching, and acceptor blinking can all produce similar changes in the apparent FRET efficiency during a burst, making them difficult to distinguish from one another (**Figure 1d**). FENNEC addresses this by operating directly on the per-photon arrival times and simultaneously classifying each burst for all three phenomena. The model is trained entirely on simulated bursts generated using parameters calibrated to the user’s instrument and sample (**Figure 1e**). Because ground-truth labels for dynamics, photobleaching, and blinking are not known in real experimental data, simulation-based training provides access to large, diverse datasets with known labels.

**Figure 1.**
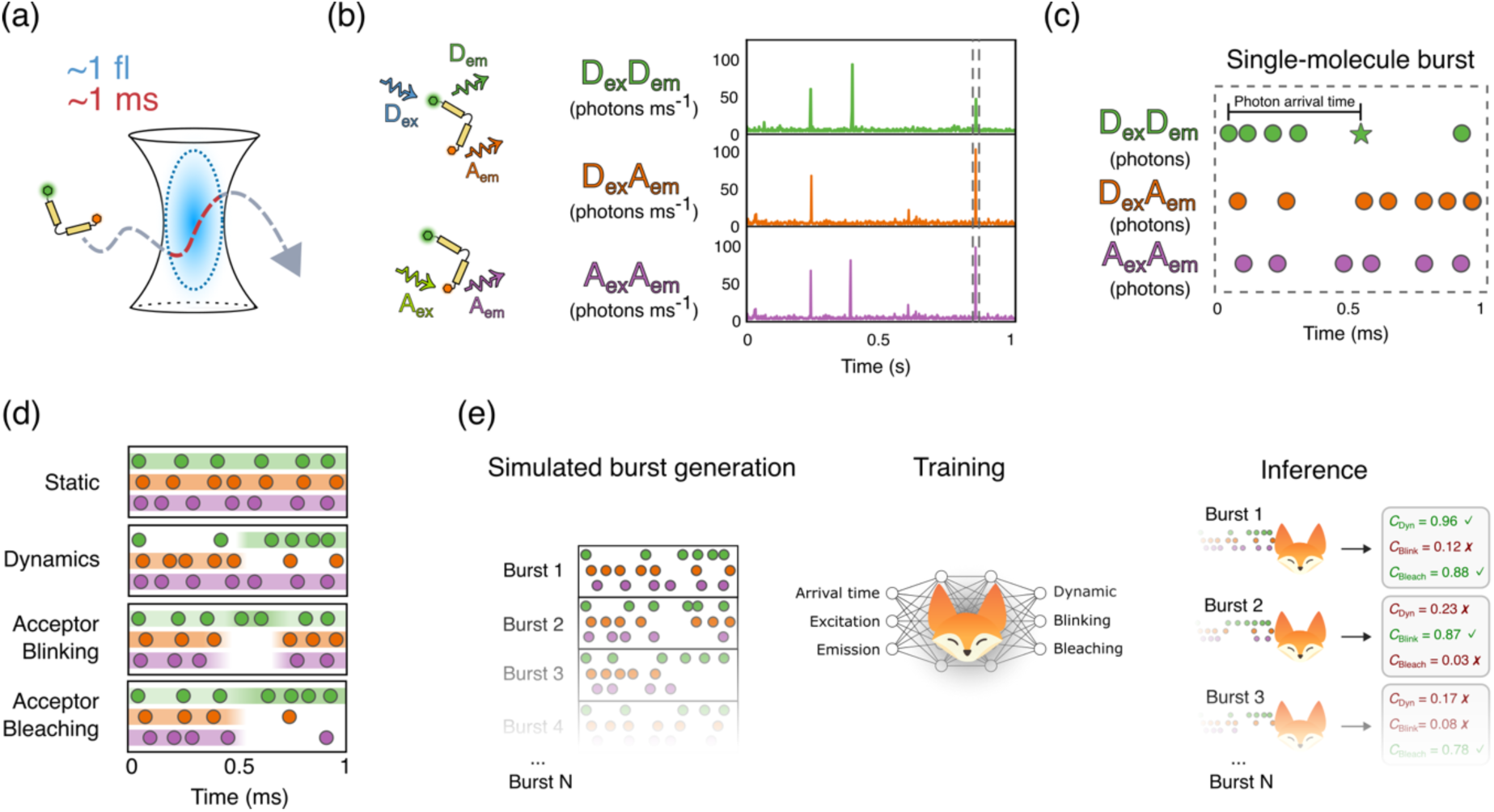
Overview of the FENNEC workflow. (**a**) Diffusion-based smFRET: freely diffusing fluorescently labelled molecules traverse a confocal excitation volume (∼1 fL), producing fluorescence bursts (∼1 ms). (**b**) ALEX and PIE yield three utilised photon streams defined by excitation and emission channel (D_ex_D_em_, D_ex_A_em_ and A_ex_A_em_; A_ex_D_em_ is recorded but not used for analysis), shown here as binned photon rates. A single-molecule burst is highlighted (dashed box). (**c**) A burst viewed at the single-photon level, showing the raw photon arrival times that serve as input to FENNEC. For the photon denoted as a star, the photon arrival time (time since the first photon) is indicated. (**d**) Four representative burst scenarios. A static molecule maintains a constant apparent FRET efficiency, whereas three main distinct phenomena produce changes during a burst: dynamics, acceptor blinking and acceptor photobleaching. The shaded background indicates the underlying emission rate in each channel (the ground truth), from which the individual photons shown are stochastic samples. (**e**) The FENNEC workflow. Simulated bursts are generated using parameters calibrated to the user’s instrument and used to train a dilated CNN. At inference, the trained model assigns each experimental burst three independent confidence scores between 0 and 1, quantifying the model’s confidence that the burst exhibits FRET dynamics (*C*_dyn_), acceptor blinking (*C*_blink_) and acceptor photobleaching (*C*_bleach_). Data shown in this figure are schematic representations for illustrative purposes and are not drawn from experimental measurements.

To approximate the experimental conditions in the simulated training data, key parameters are extracted from the user’s experimental data and used to calibrate the burst simulator (**Supplementary Figure 1**). These parameters are the burst duration distribution (**Supplementary Figure 1a**), the distribution of D_ex_D_em_ + D_ex_A_em_ emission rates (total emission under donor excitation) (**Supplementary Figure 1b**), the distribution of the A_ex_A_em_ emission rate (**Supplementary Figure 1c**), the correlation between emission rates (brightness) in each channel (**Supplementary Figure 1d**), the background rate in each channel (**Supplementary Figure 1e**), and the intensity profiles of the donor and acceptor excitation volume (**Supplementary Figure 1f**). The intensity profile is shaped by the point spread function (PSF), which determines how the detected photon rate rises and falls during the burst as the molecule diffuses through the confocal volume (Hagai & Lerner, 2019). We estimated the PSF profile separately for the donor and acceptor channels using donor-only and acceptor- only bursts, respectively, and applied these profiles to the simulated bursts.

Using these parameters calibrated to experimental data, we computationally generated millions of simulated bursts with known ground-truth labels for conformational dynamics, acceptor photobleaching, and acceptor blinking. This approach allowed us to sample a diverse combination of dynamics and FRET states that would not be achievable with experimentally labelled data.

### FENNEC classifies simulated bursts with known conformational dynamics and photophysical artefacts

We first trained FENNEC on simulated bursts labelled for static or dynamic behaviour, acceptor blinking, and acceptor bleaching (**Figure 2a-e**). Combinations of conformational dynamics and photophysical artefacts were included (see **Methods**). We then evaluated whether the trained dilated CNN could classify these simulated bursts directly from raw photon arrival times. To illustrate the challenge faced by the classifier we first visualised the training data using two common FRET data representations, efficiency-stoichiometry (E-S) and BVA plots (**Figure 2a-e**). In **Figure 2a** all simulated bursts are shown, while **Figure 2b-e** show bursts that belong exclusively to the labelled class, with co-occurring labels omitted so that its characteristic signature is visible. The top row shows the marginal FRET efficiency (E) histogram for each class. In the E-S plots (middle row), bursts dominated by donor-only emission, either from singly labelled molecules lacking an acceptor fluorophore or from molecules whose acceptor blinks or bleaches during the burst, are shifted to high S and low E. Bursts from molecules with emitting donor and acceptor fluorophores fall at intermediate S, and span a range of E values. The lower row shows BVA, which plots the within-burst standard deviation of E versus E. The black line marks the expected standard deviation for static bursts from photon shot-noise alone. Static bursts (**Figure 2b**) scatter along this line, whereas dynamic bursts (**Figure 2c**) lie above it, consistent with increased within-burst E variance due to transitions between FRET states.

**Figure 2.**
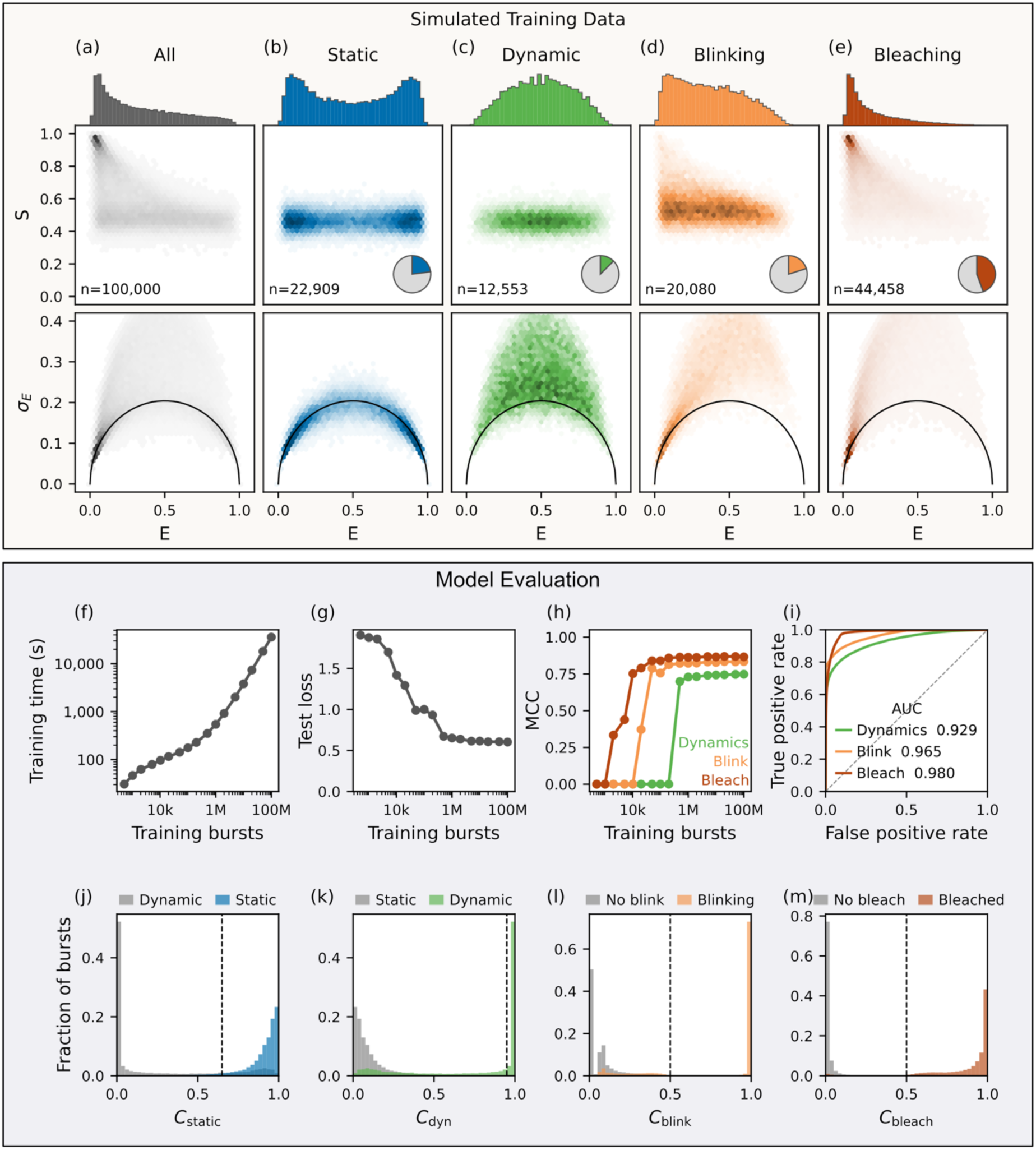
Simulated training data and FENNEC performance on data with known ground truth. (**a-e**) Composition of the simulated training data. Each column in (**b-e**) shows bursts belonging exclusively to one class, so that the characteristic signature of that class is visible without contamination from co-occurring labels (e.g. bursts that are simultaneously dynamic and blinking are omitted): (**a**) all bursts (grey), (**b**) static (blue), (**c**) dynamic (green), (**d**) blinking (orange), (**e**) bleaching (dark red). Top row, marginal FRET efficiency (E) histogram; middle row, stoichiometry (S) versus E density; bottom row, within-burst standard deviation of E (σ_E_) versus E, with the black line marking the shot-noise limit expected for static bursts (BVA). BVA was calculated with a photon window size of 6. (**f, g**) Learning curves showing (**f**) training time and (**g**) test loss versus training set size. (h) Per-task Matthews correlation coefficient (MCC) versus training set size. (**i**) ROC curves on the test set, with AUC per task. (**j-m**) Model confidence score distributions per task. Ground-truth negative classes are shown in grey and ground-truth positive classes are shown coloured: (**j**) *C*_static_, static (blue) versus dynamic (grey), where *C*_static_ = 1 − *C*_dyn_; (**k**) *C*_dyn_, dynamic (green) versus static (grey); (**l**) *C*_blink_, blinking (orange) versus no blink (grey); (m) *C*_bleach_, bleached (red) versus no bleach (grey). Dashed vertical lines mark the confidence thresholds used for filtering and classification.

Importantly, photophysical artefacts can mimic signatures associated with conformational dynamics in E-S and BVA plots. Acceptor blinking (**Figure 2d**) and bleaching (**Figure 2e**) both introduce donor-only periods within bursts, shifting density towards high S and low E. Although obvious donor-only bursts can be removed with the common practice of gating S values between ∼0.4 and ∼0.6 (Kapanidis et al., 2004), the dynamic, blinking, and bleaching populations still overlap within this region. Likewise, in the BVA plots, bursts with acceptor blinking and bleaching resemble dynamic bursts as they also increase within-burst variability above the shot-noise line. Therefore, standard E-S and BVA analyses cannot separate genuine dynamics from these artefacts, which is the ambiguity FENNEC is trained to resolve.

To evaluate FENNEC and select an appropriate training set size, we trained the network on progressively larger sets of simulated bursts and measured classification performance on a test set with known labels (**Figure 2f-i**). As the number of training bursts increased, training time grew steadily (**Figure 2f**), whereas classification performance on the test set improved rapidly before saturating at ∼10,000,000 bursts (**Figure 2g**). Per-task performance, measured by the Matthews correlation coefficient, improved with training set size (**Figure 2h**), but the amount of training data needed for improvement varied per task: bleaching and blinking plateaued after relatively few bursts (∼100,000), while dynamics needed around two orders of magnitude more data before plateauing (∼10,000,000 bursts). The model showed strong discrimination across all tasks, with areas under the receiver operating characteristic (ROC) curve of 0.929 for dynamics, 0.965 for blinking and 0.980 for bleaching (**Figure 2i**).

FENNEC outputs per-task confidence scores (*C*) for each task: *C*_dyn_, *C*_blink_ and *C*_bleach_ (**Figure 1e**). We also define *C*_static_ as 1-*C*_dyn_. These scores showed substantial separation by ground-truth class (**Figure 2j-m)**. For each task, bursts of the ground-truth positive class were assigned confidence scores concentrated near 1, while those of the negative class clustered near 0, with only a minority of bursts falling in the ambiguous range between (**Figure 2j-m**). Consistent with the ROC analysis (**Figure 2i**), this separation was cleanest for blinking and bleaching (**Figure 2l,m**) and least distinct for dynamics (**Figure 2j,k**), where a larger fraction of bursts occupied the region of overlap between the two distributions. The dashed lines mark the classification thresholds subsequently applied to the test data. Bursts with *C*_blink_ ≥ 0.5 or *C*_bleach_ ≥ 0.5 were classified as containing a photophysical artefact and excluded before static/dynamic assignment. Of the remaining bursts, those with *C*_dyn_ ≤ 0.35 were assigned as static, those with *C*_dyn_ ≥ 0.95 were assigned as dynamic, and those with intermediate scores were classified as ambiguous. These thresholds define the filtering and classification applied to all data analysed here and are user adjustable. After artefact filtering, static and dynamic assignments had precisions of 0.911 and 0.994 and recalls of 0.922 and 0.643, respectively. The stringent dynamic threshold was chosen to prioritise precision over recall for dynamic assignments. Complete performance statistics are provided in **Supplementary Table 1**.

We next benchmarked FENNEC against mpH^2^MM, an established method for resolving dynamics in diffusion-based smFRET data (Harris et al., 2022; Pirchi et al., 2016). We compared them using three simulated datasets of increasing difficulty, each with an artefact load of ∼55% blinking or bleaching bursts: a static mixture, two-state dynamics, and two-state dynamics with a separate static population at intermediate FRET (**Supplementary Figure 2a-c**). Because mpH^2^MM does not distinguish blinking from bleaching, we merged FENNEC’s blink and bleach assignments into a single artefact class to match its output resolution, and we supplied mpH^2^MM with the correct number of states for each dataset. Both methods performed comparably on the two simpler datasets, recovering the ground-truth class distributions with per-class precisions within a few percent of each other (**Supplementary Figure 2d, e**). On the hardest dataset, FENNEC outperformed mpH^2^MM in precision on every burst class, most clearly for dynamic bursts (0.96 versus 0.72 for mpH^2^MM). In this case, mpH^2^MM confused the intermediate static species with dynamic bursts that time-average to the same efficiency (**Supplementary Figure 2f**). This comparison gives mpH^2^MM an advantage, since it was supplied with the correct number of states in advance. When state number selection was performed automatically instead, mpH^2^MM overcounted under both selection criteria in the hardest condition, and under ICL in all three (**Supplementary Table 2**). In experimental data, where the true number of states is unknown, such misclassification would propagate into downstream analysis, producing spurious conformational states and unreliable transition rates.

Given the correct number of states, mpH^2^MM accurately recovered the underlying transition rates, returning 759 and 254 s^-1^ for the dynamic dataset against ground-truth values of 750 and 250 s^-1^ and near-zero exchange for the static mixture (**Supplementary Table 3**). This kinetic quantification is not provided by FENNEC. However, for the dataset containing a dynamic population mixed with a static intermediate-FRET species, the forced ground-truth four-state fit failed to resolve the true FRET states, and so did not recover meaningful rates (**Supplementary Table 3**).

Taken together, these results show that FENNEC can learn from raw, unbinned photon streams to classify dynamics, blinking and bleaching with high precision. Compared with mpH^2^MM, FENNEC performs competitively on simple cases and more strongly on the hardest, while requiring no assumption about the number of states. This positions FENNEC as a fast analysis and filtering method that can isolate clean burst populations for interpretation and downstream analysis.

### FENNEC recovers expected populations in experimental data

We next applied FENNEC to experimental data for a construct with well characterised dynamic behaviour: a DNA hairpin that interconverts between open and closed states (**Figure 3a**) (Harris et al., 2022; Tomov et al., 2012; Tsukanov et al., 2013). The per-task confidence score distributions for the experimental data closely resembled those obtained on the simulated test set (**Figure 3b**, **Figure 2j-m**), suggesting that the experimental bursts were well represented by the training simulations.

**Figure 3.**
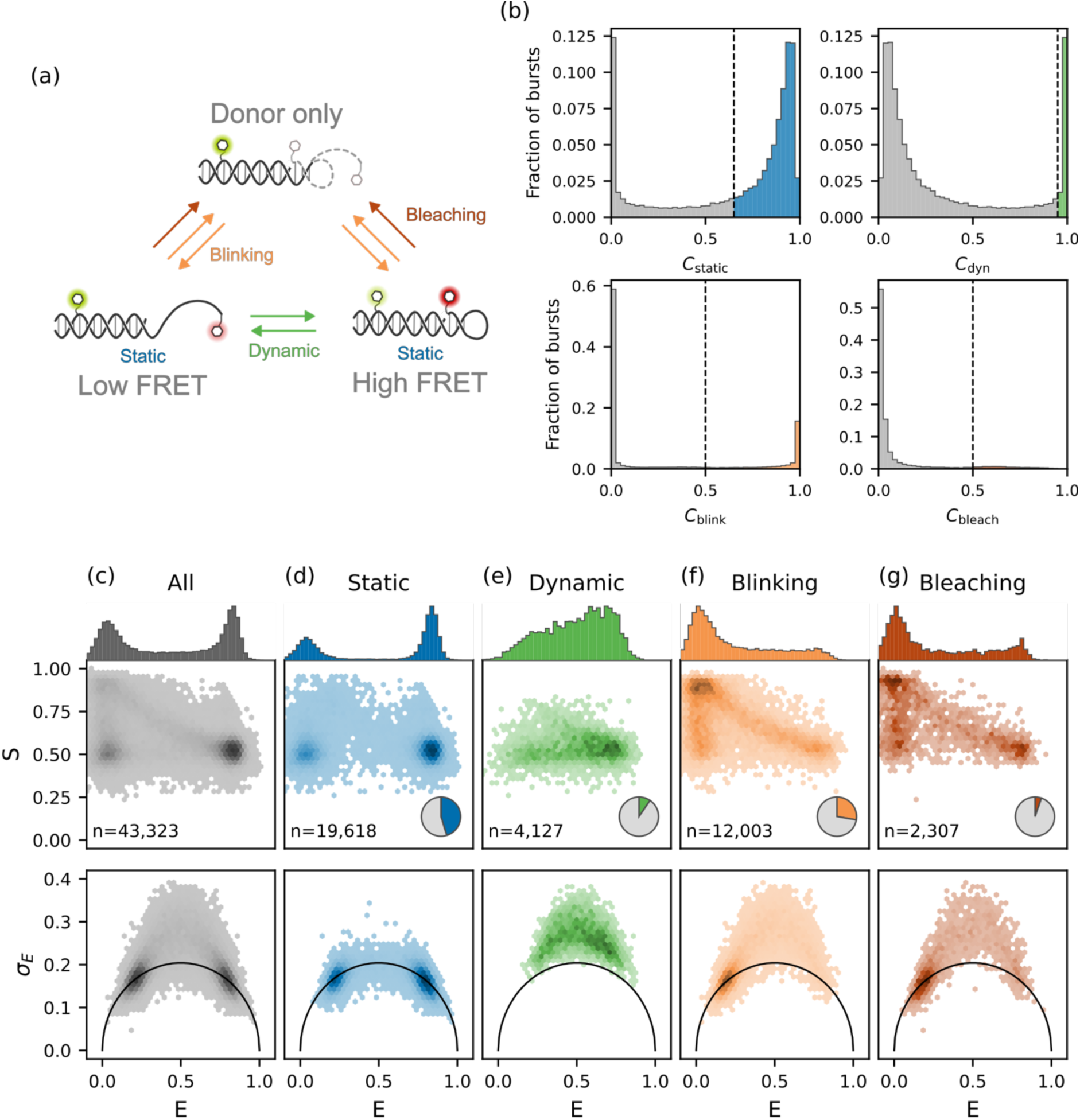
FENNEC classification of experimental bursts for a DNA hairpin reproduces expected population distributions**. (a)** Schematic of the DNA hairpin construct and the burst categories FENNEC distinguishes: static (a single FRET state), dynamic (conformational exchange between low- and high-FRET states), and the photophysical artefacts of acceptor blinking and bleaching that result in donor-only emission. **(b)** Distributions of FENNEC confidence scores across all bursts for each classification task: *C*_static_, *C*_dyn_, *C*_blink_ and *C*_bleach_. The dashed line in each panel marks the classification threshold. Bursts exceeding it (coloured) are assigned to the corresponding population. **(c-g)** Burst populations shown for **(c)** all bursts (grey) and the FENNEC-assigned **(d)** static bursts (blue), (e) dynamic bursts (green), **(f)** bursts containing blinking (orange) and **(g)** bursts containing bleaching (red). In each upper panel, the 2D histogram shows stoichiometry (S) versus FRET efficiency (E), with the marginal E histogram above. The lower panels plot the per-burst FRET standard deviation (σ_E_) against E (BVA). BVA was calculated with a photon window size of 6. Hexbin shading shows burst count per bin, scaled independently per panel, with the per-panel total bursts shown as n. Pie chart insets give each population’s fraction of all bursts.

Before classification, the DNA hairpin produced a clear bimodal E distribution, with populations at low and high E corresponding to the open and closed conformations (**Figure 3c**, upper). The corresponding BVA panel showed a broad arc of excess variance rising well above the shot-noise curve across intermediate E values (**Figure 3c**, lower), the signature conventionally taken to indicate FRET dynamics. Taken at face value, the construct therefore appears strongly dynamic. We then applied FENNEC and separated the data into different burst classes. The largest class comprised bursts assigned as static (n = 19,618; 45%) and their distribution reproduced the low and high E peaks in the E-S plots, residing on the shot-noise curve in BVA (**Figure 3d**), consistent with the expected two static open and closed states. Bursts assigned as dynamic formed a smaller population (n = 4,127; 10%) at intermediate E with the expected stoichiometry (S ≈ 0.5), bridging the two static states and lying clearly above the BVA shot-noise curve (**Figure 3e**). This is the expected signature of molecules interconverting on timescales equal to or shorter than the burst duration. Importantly, this genuine dynamic population accounted for only a small fraction of the excess variance in the all-burst panel.

FENNEC attributed most of the intermediate-E and above-curve BVA signal to acceptor blinking and bleaching (**Figure 3f, g**). These bursts follow a diagonal trend in E-S space, apparent E decreasing and S increasing as acceptor signal is lost. It is this diagonal smear, rather than conformational exchange, that generates much of the apparent low-FRET arc and the elevated BVA variance. Blinking accounted for 28% of bursts (n = 12,003), substantially more than bleaching (n = 2,307; 5%). Without per-burst classification this artefact signal overlaps the genuine dynamic region in E, S and BVA space and would be difficult to remove with high precision. FENNEC separates it directly, leaving a clean dynamic population for interpretation and downstream analysis.

Finally, we compared FENNEC’s burst classifications directly against mpH^2^MM on the same experimental data (**Supplementary Figure 3**). The two methods agreed on 86.8% of FENNEC-classified bursts (**Supplementary Figure 4**). We inspected the remaining classification calls (13.2%) where the methods disagreed using E-S and BVA plots to evaluate the assignments (**Supplementary Figure 5**). In calls containing artefacts, neither method was clearly better supported. However, where the methods disagreed on static-versus-dynamic calls (in either direction), BVA supported FENNEC’s assignment.

## Discussion

Deep learning approaches are increasingly being used in analysing time-resolved and spectroscopic biophysical data (Beigzadeh et al., 2026). In surface-immobilised smFRET, methods such as DeepFRET (Thomsen et al., 2020), Deep-LASI (Wanninger et al., 2023), and META-SiM (Li et al., 2025) now automate classification and segmentation of time traces, tasks that previously required extensive manual curation. Similar approaches have been developed in magnetic resonance spectroscopy using neural networks trained on simulated time-domain data: DEERNet learns from simulated DEER time traces to recover distance distributions from pulsed EPR data (Worswick et al., 2018), while FID-Net applies a WaveNet-inspired architecture directly to time-domain NMR data for spectral reconstruction and virtual decoupling (Karunanithy & Hansen, 2021; Karunanithy et al., 2021, 2024). The use of machine learning to filter artefactual data is well established in single-particle cryo-EM, where contamination from ice crystals, carbon edges, and aggregates must be removed to avoid degrading downstream 3D reconstruction. Deep learning tools now automate this filtering at multiple stages of the pipeline, from CNN-based particle pickers (Bepler et al., 2019; Sanchez-Garcia et al., 2020) to automated 2D class selection (Kimanius et al., 2021). Notably, PARSED demonstrated that training on purely simulated data can achieve competitive picking performance without experimental labels (Yao et al., 2020), an approach conceptually analogous to FENNEC.

Several features distinguish FENNEC from existing burst classification approaches. Its central practical advantage is that classification requires no prior assumption about the number of underlying conformational states, in contrast to HMM-based methods, which require candidate state numbers to be specified and compared before interpretation. This positions FENNEC as a first filter when the system is poorly characterised, and the per-burst assignments it returns, each with an associated confidence, provide an immediate triage of a dataset rather than a single global kinetic model. The practical value of this filtering is evident in the experimental data, where ∼33% of bursts showed blinking and bleaching, a large fraction that would otherwise distort any quantification of conformational exchange.

FENNEC shows that a deep neural network operating directly on raw photon arrival times can separate conformational dynamics from photophysical artefacts in diffusion-based smFRET. It provides a fast, per-burst analysis that complements established methods. Its outputs are straightforward to interpret, turning the raw photon arrival times in each burst into an immediate overview of the static, dynamic, and artefactual components of a dataset. We demonstrate FENNEC on a well-characterised DNA construct as a proof of principle, and release it for open use by the smFRET community.

## Methods

### Single-molecule FRET sample preparation

Oligonucleotides were purchased from Integrated DNA Technologies (IDT) as lyophilised solids and reconstituted in DEPC-treated water to stock concentrations of 100 μM. Fluorophores were conjugated to either the 5’ end or an internal amino-modified thymidine; the labelled position is marked in each sequence by asterisks flanking the modified base (*T*). The DNA hairpin was assembled from an 80-base strand (5’-TCG CCA AAA AAA AAA AAA AAA AAA AAA AAA AAA AAA ATG GCG ATT TTC TTC ACA AAC CAG TCC AAA CTA TCA CAA ACT TA-3’) labelled at the 5’ end with ATTO 647N (ATTO-TEC), and a 38-base strand (5’-TTT TTA AGT TTG TGA TAG TTT GGA CTG G*T*T GTG AAG AA-3’) labelled internally with ATTO 565 (ATTO-TEC) and bearing a 5’ biotin (the biotin was not used in this study). Strands were mixed at equimolar concentrations of 10 μM in hybridisation buffer (20 mM Tris-HCl pH 7.5, 150 mM NaCl, 2 mM MgCl_2_), heated to 90 °C for 3 minutes in a water bath, then cooled slowly to room temperature on the bench protected from light. Hybridised samples were stored at 4 °C for short-term use or at −80 °C for longer-term storage. Immediately before measurement, samples were diluted to approximately 50 pM in TBS (20 mM Tris-HCl pH 7.5, 150 mM NaCl).

### Single-molecule FRET data acquisition

smFRET experiments were performed on a Luminosa single-photon-counting confocal microscope (PicoQuant) equipped with an Olympus UPlanSApo 60x/1.20 NA water-immersion objective. PIE was achieved at a combined repetition rate of 40 MHz using 560 nm and 640 nm lasers with average powers of 80 μW and 40 μW at the sample, respectively. Excitation light was directed onto the sample via a DB532/640 dual-band major dichroic. Emitted fluorescence was passed back through the major dichroic, separated into donor and acceptor channels by a 635 nm long-pass dichroic, and detected on two single-photon avalanche diodes through a 582/64 nm bandpass filter (donor channel) and a 690/70 nm bandpass filter (acceptor channel). The variable PSF feature (VarPSF) was set to the largest available confocal volume and the pinhole diameter was set to 100 μm. For each measurement, 100 μl of sample at approximately 50 pM was loaded into a glass bottom 18-well μ-Slide chamber (81817, Ibidi), and the focal spot was positioned 50 μm above the coverslip surface. Data were acquired for 6 hours at room temperature (21 ± 1 °C).

### Generation of simulated training data

FENNEC is trained entirely on simulated smFRET bursts with parameters calibrated to real experimental data. Each burst is modelled as a Poisson photon arrival process across three detection channels (D_ex_D_em_, D_ex_A_em_, and A_ex_A_em_), with parameters drawn from distributions fitted to a reference dataset and includes the two acceptor photophysical processes most often confounded with conformational dynamics: blinking and photobleaching. Full details are provided in the **Supplementary Methods**.

### Model architecture and training

Each photon is represented by three features: log-transformed arrival time, excitation channel, and emission channel. Bursts are processed by a gated dilated one-dimensional convolutional network with three independent classification heads for dynamics, blinking and bleaching trained on simulated bursts. Full details are provided in the **Supplementary Methods**.

### Inference on experimental data

Experimental bursts are identified using FRETBursts (Ingargiola et al., 2016) and a dual-channel burst search (Nir et al., 2006) and featurised identically to the training data, then passed through the trained FENNEC model to yield independent confidences for the three tasks (*C*_dyn_,*C*_blink_ and *C*_bleach_), with class assignment based on per-task confidence thresholds. Full details are provided in the **Supplementary Methods**.

## Supplementary Material

Supplementary information – Document containing the Supplementary Methods, Supplementary Figures 1-5 and Supplementary Tables 1-3.

## Data Availability

The DNA hairpin dataset used in this study is available on Zenodo at https://doi.org/10.5281/zenodo.21135449 in Photon-HDF5 format. The FENNEC source code, including the simulation framework, network architecture, training pipeline and inference tools, is available at https://github.com/jacrossley/FENNEC.

## Acknowledgements

**B.S.** and **J.A.C.** acknowledge support and funding from the University of Leeds and the Astbury Centre for Structural Molecular Biology. **J.A.C.** also acknowledges the BBSRC for funding (BB/Y00034X/1). The PicoQuant Luminosa was funded through the Cheney Biomedical Accelerator, via the generous gift of Peter & Susan Cheney and the Wolfson Foundation (360G-wolfson-24625). We also thank all members of the Sheena E. Radford, David J. Brockwell and Antonio N. Calabrese labs for insightful discussions and support. This work was undertaken in part on the Aire HPC system at the University of Leeds, UK.

## Author Contributions

**B.S.**: Conceptualisation, Methodology, Software, Writing - Original Draft, Writing - Review & Editing. **J.A.C.**: Conceptualisation, Methodology, Software, Investigation, Writing - Original Draft, Writing - Review & Editing, Visualisation, Supervision, Project administration, Funding acquisition.

## Declaration Of Interests

The authors declare no competing interests.

## Notes

https://github.com/jacrossley/FENNEC

https://doi.org/10.5281/zenodo.21135449

### Competing Interest Statement

The authors have declared no competing interest.

