## Supplementary Information for "FENNEC: photon-level deep learning for classifying bursts in diffusion-based single-molecule FRET"

### Supplementary Methods

#### Generation of simulated training data

FENNEC is trained entirely on simulated smFRET data generated by a burst simulator calibrated to real experimental data. Each burst is simulated as a Poisson photon arrival process across three detection channels corresponding to the ALEX/PIE excitation scheme: donor excitation and donor emission ($D_{ex}D_{em}$), donor excitation and acceptor emission ($D_{ex}A_{em}$), and acceptor excitation and acceptor emission ($A_{ex}A_{em}$).

Burst-level parameters are drawn for each simulated burst from distributions fitted to real smFRET measurements (**Supplementary Figure 1**). PTU files from SymPhoTime64 software are converted into Photon-HDF5 data files using phconvert v0.10.1 (Ingargiola, Laurence, et al. 2016). These files are then analysed using the FRETBursts v0.8.3 Python package (Ingargiola, Lerner, et al. 2016). Single-molecule bursts were extracted from the data stream using the dual-channel burst search (DCBS) (Nir et al. 2006) and corrections applied: donor leakage ($\alpha = 0.165$), direct acceptor excitation ($\delta = 0.06$), detection efficiency ($\gamma = 0.8$), and excitation efficiency ($\beta = 1.5$).

The burst duration is fit to a log-normal distribution clipped at the 0.1 and 99.9 percentiles (**Supplementary Figure 1a**). FRET emission ($D_{ex}D_{em}$ + $D_{ex}A_{em}$) rates were taken from FRET-population bursts (DCBS & S = 0.4-0.6) in the reference dataset and fitted to a log-normal distribution clipped at the 0.1 and 99.9 percentiles (**Supplementary Figure 1b**). Acceptor emission ($A_{ex}A_{em}$) rates were taken from FRET-population bursts (DCBS & S = 0.4-0.6) in the reference dataset and fitted to a log-normal distribution clipped at the 0.1 and 99.9 percentiles (**Supplementary Figure 1c**). The total donor-excitation photon rate ($D_{ex}D_{em}$ + $D_{ex}A_{em}$) and the $A_{ex}A_{em}$ photon rate are positively correlated: bursts from molecules transiting near the focal centre are brighter in both excitation periods, while peripheral transits are faint in both

(**Supplementary Figure 1d**). To reproduce this coupling, in the burst simulator the two rates are drawn jointly from a bivariate log-normal distribution rather than independently. Pairs (log $R_{Dex}$, log $R_{Aex}$) are sampled from this bivariate normal and exponentiated to give per-burst rates. In the burst generator donor-excitation photons are partitioned between the $D_{ex}D_{em}$ and $D_{ex}A_{em}$ channels according to the efficiency of the current FRET state, such that the $D_{ex}A_{em}$ rate equals the total donor-excitation rate multiplied by E and the $D_{ex}D_{em}$ rate equals the remainder.

Background photon rates were estimated separately for each channel ($D_{ex}D_{em}$, $D_{ex}A_{em}$ and $A_{ex}A_{em}$) using FRETBursts (**Supplementary Figure 1e**). These per-channel rates were then added to every simulated burst as an independent Poisson process.

To reproduce the spatial intensity variation encountered in confocal detection, burst rates are modulated by an empirical intensity profile extracted from single-label bursts in the reference dataset using a single-channel burst search (**Supplementary Figure 1f**): the $D_{ex}D_{em}$ profile from donor-only bursts ($S > 0.9$, $E < 0.15$) and the $A_{ex}A_{em}$ profile from acceptor-only bursts ($S < 0.2$). Each burst is divided into 20 equal-width positional bins spanning its duration, and the mean photon rate per bin (normalised to the burst-averaged rate) is used as a multiplicative envelope applied to the corresponding signal channel during simulation. Background contributions are not modulated.

Two acceptor photophysical processes are modelled explicitly. Acceptor photobleaching is treated as an irreversible exponential process with a rate constant $k_{bleach} = 80\ s^{-1}$; following a bleaching event, the $D_{ex}A_{em}$ and $A_{ex}A_{em}$ photon rates drop to their respective background levels while the $D_{ex}D_{em}$ rate rises to the full donor-excitation rate. Acceptor blinking into a non-fluorescent triplet state is modelled with an intersystem crossing yield $\Phi_{ISC} = 0.002$. Acceptor blink events occur as a Poisson process with rate $k_{blink} = (R_{Dex} + R_{Aex})\cdot\Phi_{ISC}$, proportional to the burst's total photon rate, where $R_{Dex}$ is the donor-excitation rate (the combined $D_{ex}D_{em} + D_{ex}A_{em}$ rate) and $R_{Aex}$ is the acceptor-excitation rate; both are the signal rates sampled for the burst, before application of the PSF envelope and background. Dark-state dwell times are drawn from an exponential distribution with mean $\tau_{triplet} = 1$ ms. The minimum resolvable dark-state duration is 500 μs; any burst containing a sub-threshold blink is discarded entirely to avoid label ambiguity.

Fixed per-channel background rates, measured from inter-burst regions of real experimental data, are added to all simulated bursts. Simulated bursts containing fewer than 100 donor-excitation photons ($D_{ex}D_{em}$ + $D_{ex}A_{em}$) were discarded to match the minimum burst size criterion applied during experimental burst search. Bursts were additionally filtered to match the sensitivity threshold of the DCBS used on real data, applied to the donor-excitation period only: a burst was rejected and resampled unless its donor-excitation rate plus the summed donor-excitation background exceeded $F = 6$ times that background. This ensured the training distribution contained only bursts detectable under the same criteria applied to experimental data. Bursts exceeding 1000 total photons were discarded to maintain a fixed input length for the network (~2.5% of bursts in the DNA Hairpin dataset). We discarded bursts containing very short blink events (< 500 μs) or state dwell times (< 100 μs), as these fall below the temporal resolution the model can reasonably learn. We also removed bursts in which FRET transitions occurred only while the acceptor was dark, whether through blinking or after bleaching, since the dynamics leave no detectable signature when the acceptor is not emitting and the model cannot fundamentally learn to identify them.

**Model architecture and training**

Each photon in a burst is represented by three features: the log-transformed time since the first photon in the burst (log1p, in μs), the excitation channel (0 for donor, 1 for acceptor), and the emission channel (0 for donor, 1 for acceptor). The log1p transformation compresses the dynamic range of photon timing, which spans several orders of magnitude within a single burst. A log1p transformation, $\log(x + 1)$, rather than log is used to ensure the zero timepoint is defined. All bursts are padded to a fixed length of 1000 photons or discarded if they are longer, and a Boolean padding mask is propagated to all downstream layers to prevent padded positions from contributing to learned representations.

The network is a gated dilated one-dimensional convolutional neural network operating on the per-photon feature sequence. A linear layer first projects the three input features to 64 channels. The core comprises nine gated dilated convolutional blocks with kernel size 3 and exponentially increasing dilations (1, 2, 4, 8, 16, 32, 64, 128, 256). Each block applies a gated activation, the elementwise product of a tanh-activated filter convolution and a sigmoid-activated gate convolution. The gated activated output then contributes both a skip projection to 128 channels and a 1×1 residual projection added back to the block input. The skip outputs of all nine blocks are summed, giving a receptive field of 1,023 photons that spans the full 1000-photon input. The summed skip signal passes through a ReLU, a 1×1 convolution (128→128), a further ReLU, and a second 1×1 convolution (128→128). A masked mean-pool over non-padded photons yields a single 128-dimensional burst-level representation. This feeds a shared linear layer (128→256) with ReLU activation, followed by three independent linear heads (256→2) producing logits for dynamics, bleaching, and blinking respectively. The complete model comprises 402,502 parameters.

Training data are generated on-the-fly using the simulation framework described above. The model in this manuscript was trained for a single epoch consisting of 100,000,000 simulated bursts, with a fixed held-out test set of 100,000 pre-generated bursts used for evaluation. The loss function is the unweighted sum of three two-class softmax cross-entropy terms, one per classification task, optimised jointly using Adam (Kingma and Ba 2014) (learning rate $1\times10^{-3}$, weight decay $1\times10^{-4}$) with a batch size of 256. On-the-fly burst generation used seven dataloader worker processes, and training was performed on a single NVIDIA L40S GPU.

**Inference on experimental data**

PTU files from SymPhoTime64 software are converted into Photon-HDF5 data files using phconvert v0.10.1 (Ingargiola, Laurence, et al. 2016). These files are then analysed using the FRETBursts v0.8.3 python package (Ingargiola, Lerner, et al. 2016). Correction factors are applied ($\alpha = 0.165$, $\delta = 0.06$, $\gamma = 0.8$, $\beta = 1.5$). These corrections affect only the calculation of E and S for visualisation and downstream analysis; the photon arrival times and channel assignments passed to FENNEC are unmodified. Background rates are estimated per 60-second window. Bursts are identified using DCBS, requiring $m = 10$ consecutive photons at a threshold of $F = 6$ times the local background rate. Bursts containing fewer than 100 donor-excitation photons ($D_{ex}D_{em} + D_{ex}A_{em}$) are discarded.

Each burst is featurised identically to the training data. Photon arrival timestamps are separated into three channels ($D_{ex}D_{em}$, $D_{ex}A_{em}$ and $A_{ex}A_{em}$) and aligned to the first photon in the burst. Each photon is then encoded as three features: log-transformed time since burst start (log1p, in μs), excitation channel (0 for donor, 1 for acceptor), and emission channel (0 for donor, 1 for acceptor). Bursts exceeding 1000 photons are excluded entirely rather than truncated, to avoid classifying an incomplete photon record.

Featurised bursts are passed through the trained FENNEC model in mini-batches of 512. The three classification heads produce independent softmax confidences for dynamics (static or dynamic), bleaching (clean or bleached), and blinking (clean or blinked). Class assignment uses tuned per-task confidence thresholds rather than argmax. For dynamics, a dual threshold is applied: bursts with $C_{dyn} \geq 0.95$ are classified as dynamic and those with $C_{dyn} \leq 0.35$ as static, while intermediate values are excluded as ambiguous. Because the dynamics head is a two-class softmax, the static and dynamic confidences are complementary, $C_{\text{static}} = 1 - C_{\text{dyn}}$, so the static cut-off $C_{\text{dyn}} \leq 0.35$ is equivalent to $C_{\text{static}} \geq 0.65$. For acceptor bleaching, bursts with $C_{\text{bleach}} \geq 0.50$ are classified as bleached. For acceptor blinking, bursts with $C_{\text{blink}} \geq 0.50$ are classified as blinked. These classification thresholds are user adjustable for inference on experimental data using a trained model.

### Supplementary Figures

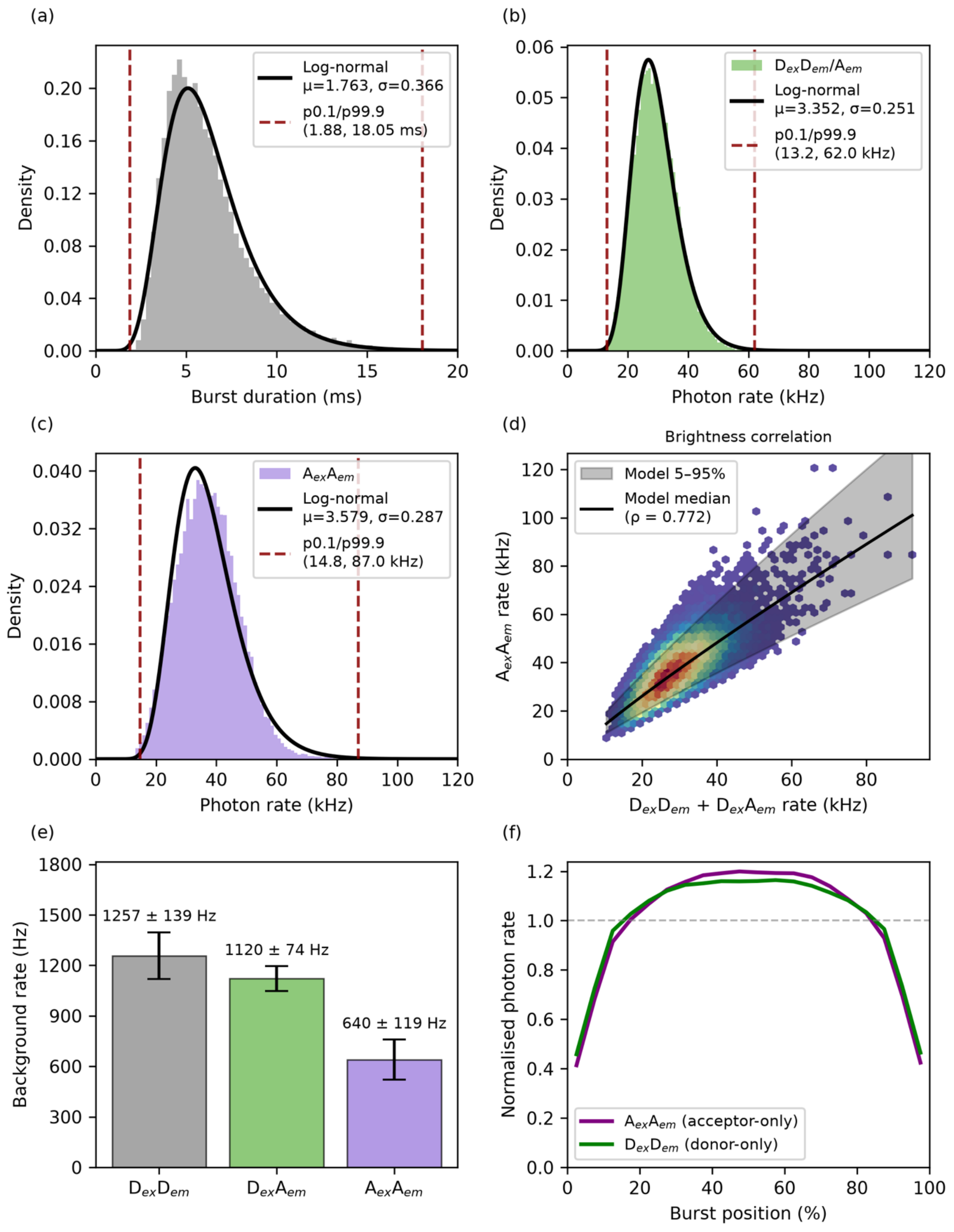


**Supplementary Figure 1.** Calibration of simulation parameters from experimental data. (**a**) Burst duration distribution fitted with a log-normal function. (**b**) Total donor-excitation photon rate ($D_{ex}D_{em}$ + $D_{ex}A_{em}$) fitted with a log-normal distribution. (**c**) Acceptor-excitation photon rate ($A_{ex}A_{em}$) fitted with a log-normal distribution. Red dashed lines in the three distribution panels mark the 0.1 and 99.9 percentiles, used as clipping bounds during burst generation. (**d**) Joint distribution of donor-excitation and acceptor-excitation rates per burst (hexbin density). The black curve shows the median of a bivariate log-normal model, and the grey band shows its 5-95% conditional interval. (**e**) Mean background photon rates in each detection stream. (**f**) Normalised photon-rate profile across the burst as a function of relative position, computed separately from $D_{ex}D_{em}$ photons in donor-only bursts ($S > 0.9$, $E < 0.15$; green) and $A_{ex}A_{em}$ photons in acceptor-only bursts ($S < 0.2$; purple).

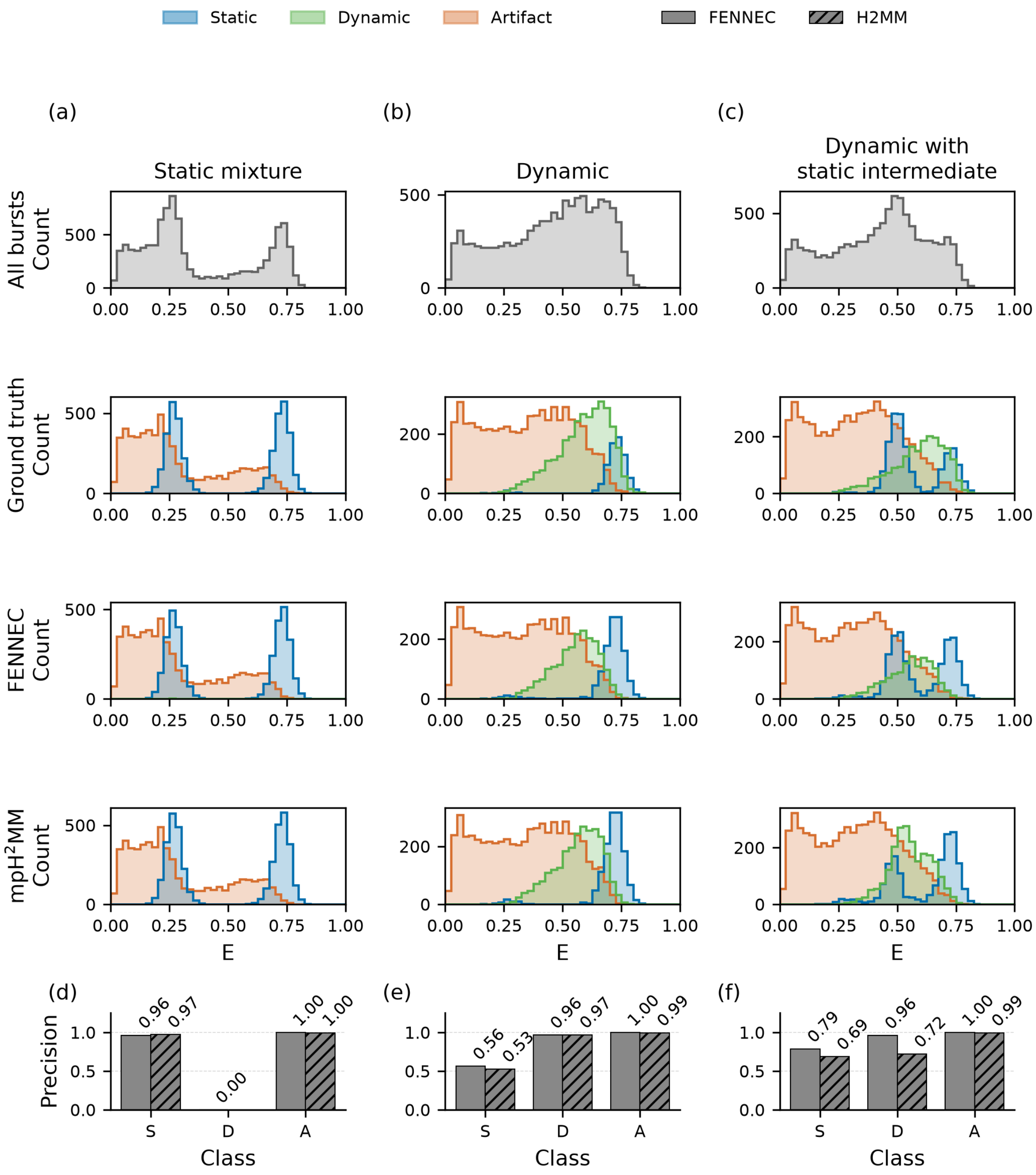


**Supplementary Figure 2.** FENNEC and mpH$^2$MM classification compared on simulated data. Each column corresponds to a simulated dataset of 10,000 bursts, comparable to the number typically obtained in smFRET experiments. (**a**) a static mixture of two non-exchanging species at E = 0.25 and 0.75 (50:50); (**b**) a two-state population exchanging between E = 0.25 and 0.75 within bursts (forward and reverse rates of 750 s$^{-1}$ and 250 s$^{-1}$, respectively); and (**c**) a 70:30 mixture of that dynamic population and a separate, non-exchanging static species at the intermediate value E = 0.5. In all three, bursts showing blinking or bleaching are grouped as a single artefact class for comparison with mpH$^2$MM, which does not distinguish between the two. Within each column, the top row is the FRET efficiency (E) histogram of all bursts (grey), and the three rows below show the same bursts split into static (blue), dynamic (green) and artefact (orange) by the ground-truth labels, the FENNEC classification and the mpH$^2$MM classification respectively. (**d-f**) Per-class precision against ground truth for FENNEC (solid bars) and mpH$^2$MM (hatched bars), for the static (S), dynamic (D) and artefact (A) classes, with values printed above each bar. mpH$^2$MM was supplied with the ground-truth number of states (three for (a) and (b), four for (c), in each case the FRET states plus one donor-only state).

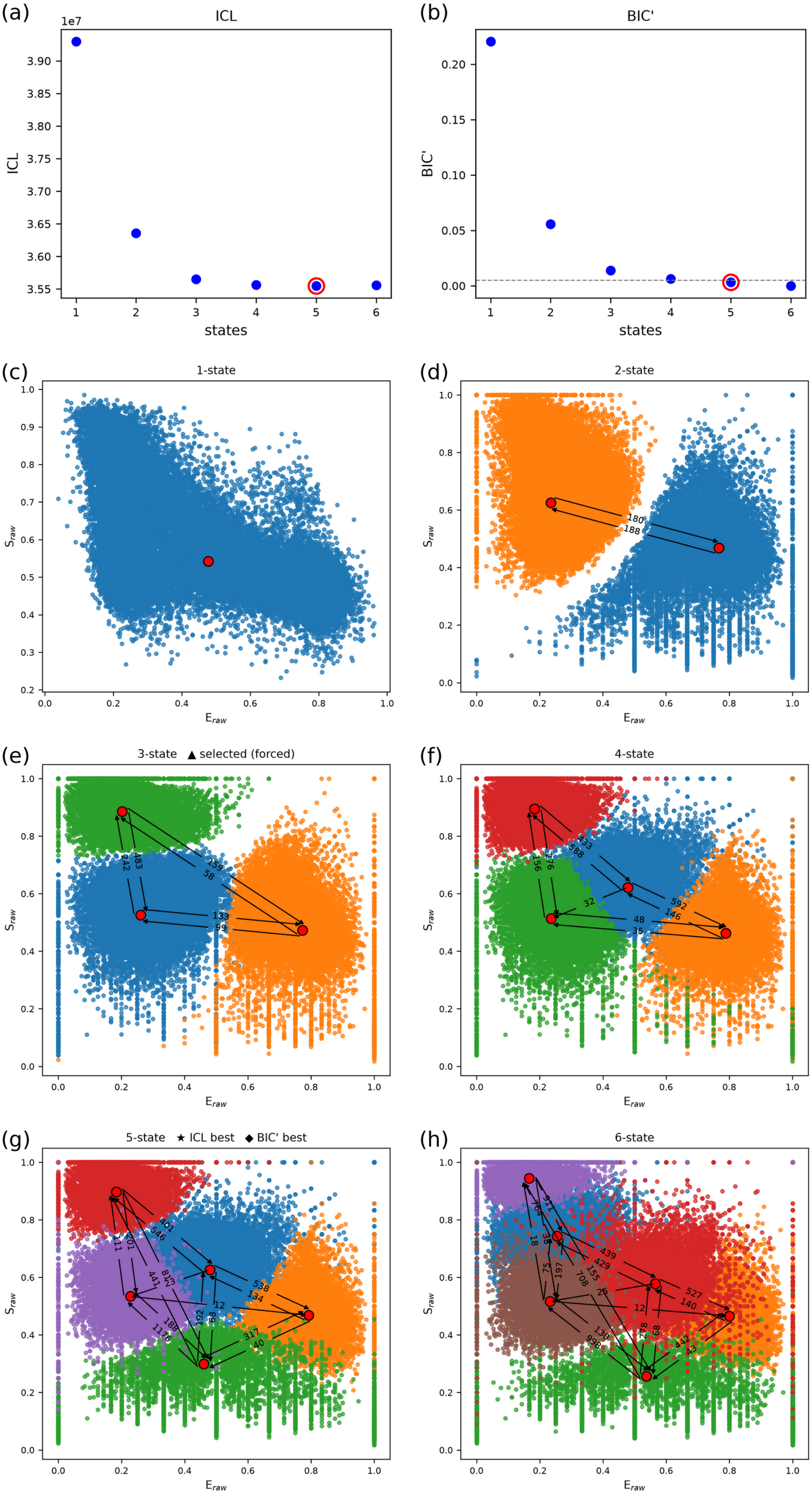

(a)
ICL
1e7
ICL
states
(b)
BIC'
BIC'
states
(c)
1-state
(d)
2-state
180
188
(e)
3-state ▲ selected (forced)
(f)
4-state
(g)
5-state ★ ICL best ◆ BIC' best
(h)
6-state
$S_{raw}$
$E_{raw}$

**Supplementary Figure 3.** mpH$^2$MM model selection and dwell-time E-S plots for the DNA hairpin dataset. (**a**) ICL and (**b**) BIC' as a function of the number of mpH$^2$MM states. Red circles highlight the states chosen by each criterion. (**c-h**) E-S dwell plots for the (**c**) one-, (**d**) two-, (**e**) three-, (**f**) four-, (**g**) five-, (**h**) six-state models, with each dwell coloured by state. Red markers indicate the fitted state positions and arrows denote transitions between states. The numbers are the fitted rate constants between states ($s^{-1}$).

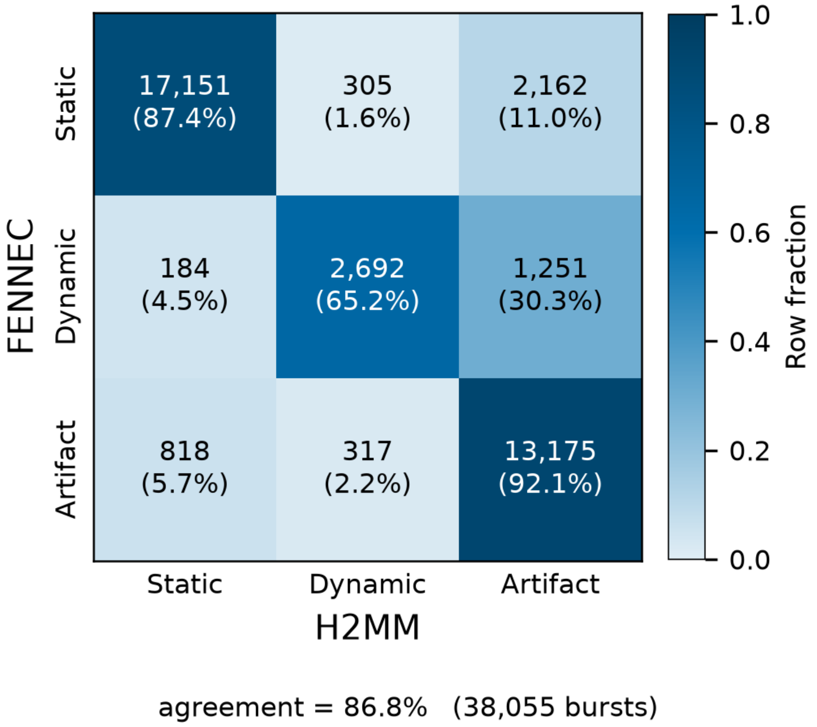


**Supplementary Figure 4.** Burst-level classification agreement between FENNEC and mpH$^2$MM on the DNA Hairpin dataset. Confusion matrix comparing FENNEC's burst classification (static, dynamic, artefact (blinking or bleaching); rows) against mpH$^2$MM (columns). Each cell reports the number of bursts and, in parentheses, the corresponding row fraction, i.e. the proportion of bursts assigned to a given FENNEC class that fall into each mpH$^2$MM class; cell colour encodes this row fraction (colour bar on the right). As neither method provides ground truth on experimental data, the matrix reports classification concordance rather than accuracy.

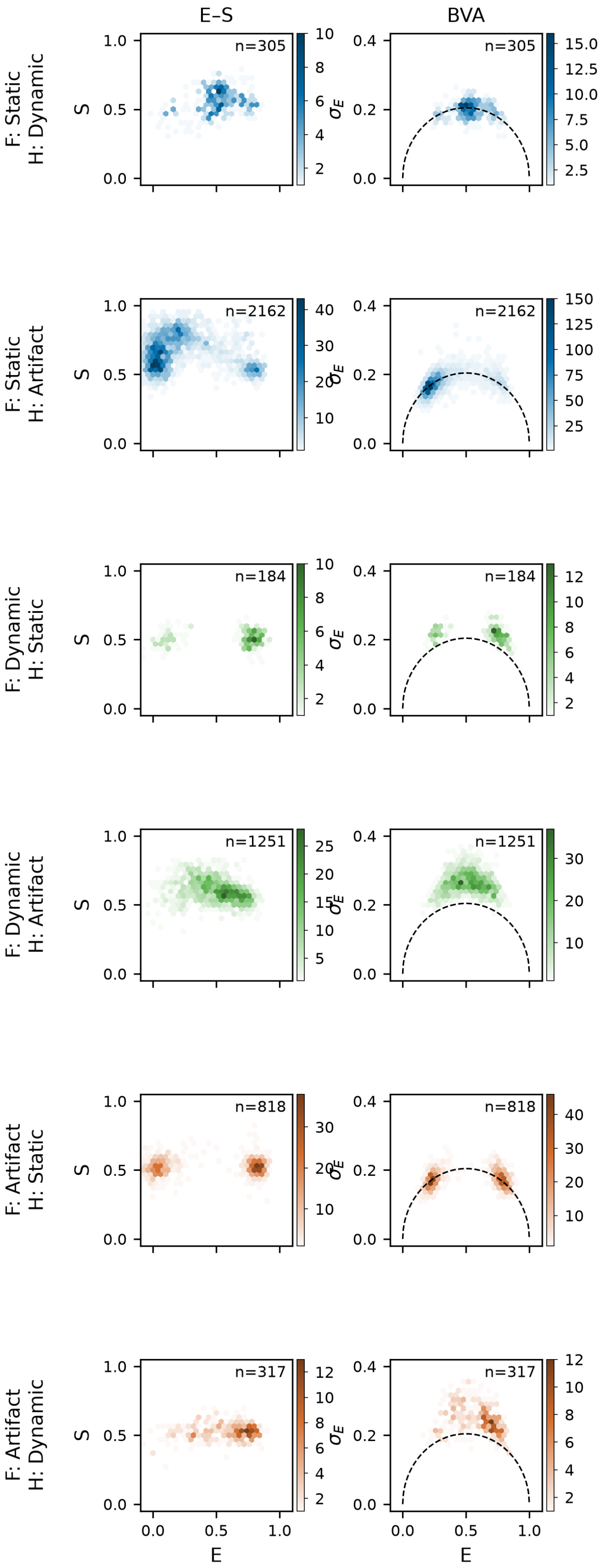
E–S
BVA
F: Static
H: Dynamic
F: Static
H: Artifact
F: Dynamic
H: Static
F: Dynamic
H: Artifact
F: Artifact
H: Static
F: Artifact
H: Dynamic
n=305
n=2162
n=184
n=1251
n=818
n=317
S
$\sigma_E$
E

**Supplementary Figure 5.** E–S and BVA plots for FENNEC / mpH$^2$MM disagreement cases. Plots for the off-diagonal cells of the confusion matrices in **Supplementary Figure 4**, i.e. the bursts on which FENNEC and mpH$^2$MM assign different classes. Each row corresponds to one disagreement category, labelled by the FENNEC call and the conflicting mpH$^2$MM call. The left column shows the E-S plot and the right column the BVA plot. Marker colour encodes the FENNEC class (blue, static; green, dynamic; orange, artefact (acceptor blinking or acceptor bleaching)), and the number of bursts per panel is indicated.

### Supplementary Tables

| Classification | Threshold | Precision | Recall | MCC |
|---|---|---|---|---|
| Static | $C_{\mathrm{dyn}} \leq 0.35$ | 0.911 | 0.922 | 0.761 |
| Dynamic | $C_{\mathrm{dyn}} \geq 0.95$ | 0.994 | 0.643 | 0.729 |
| Blinking | $C_{\mathrm{blink}} \geq 0.5$ | 0.993 | 0.766 | 0.833 |
| Bleaching | $C_{\mathrm{bleach}} \geq 0.5$ | 0.899 | 0.955 | 0.866 |

**Supplementary Table 1. FENNEC threshold-dependent classification performance on simulated test data.** Static and dynamic performance was evaluated among bursts remaining after excluding those classified as blinking or bleaching. Bursts with intermediate dynamics scores were classified as ambiguous and contributed to false negatives when calculating recall. Blinking and bleaching were evaluated independently on the complete test set. Dynamics, blinking and bleaching were treated as independent labels and could co-occur within a burst. ROC-AUC values are shown in main text **Figure 2i**.

| Condition | Ground truth | ICL | BIC’ |
|---|---|---|---|
| Static mixture | 3 | 4 | 3 |
| Dynamic | 3 | 5 | 3 |
| Dynamic with static intermediate | 4 | 6 | 5 |

**Supplementary Table 2.** State-number selection on the three benchmark datasets. For each simulated dataset, the ground-truth number of states (the FRET states plus one donor-only state) is compared with the number selected using two model-selection criteria, the integrated completed likelihood (ICL) and the modified Bayesian information criterion (BIC′). For ICL the lowest value is taken and for BIC’ the first value below 0.005 is taken as the number of states. Classification elsewhere in this work used the ground-truth state count rather than a selected one.

| Condition | State | E | S | Assignment | k to S0 $s^{-1}$ | k to S1 $s^{-1}$ | k to S2 $s^{-1}$ | k to S3 $s^{-1}$ |
|---|---|---|---|---|---|---|---|---|
| Static mixture | S0 | 0.73 | 0.45 | High-FRET | | 0 | 158.19 | |
| | S1 | 0.27 | 0.45 | Low-FRET | 0 | | 155.29 | |
| | S2 | 0.04 | 0.98 | Donor-only | 82.30 | 77.81 | | |
| Dynamic | S0 | 0.73 | 0.45 | High-FRET | | 253.89 | 157.19 | |
| | S1 | 0.27 | 0.45 | Low-FRET | 758.88 | | 172.56 | |
| | S2 | 0.04 | 0.98 | Donor-only | 114.71 | 50.99 | | |
| Dynamic with static intermediate | S0 | 0.72 | 0.50 | High-FRET | | 54.92 | 402.42 | 160.72 |
| | S1 | 0.71 | 0.39 | Mid-FRET | 12.96 | | 371.93 | 146.39 |
| | S2 | 0.37 | 0.45 | Low-FRET | 327.69 | 278.69 | | 166.32 |
| | S3 | 0.04 | 0.98 | Donor-only | 50.59 | 44.72 | 75.76 | |

**Supplementary Table 3.** mpH$^2$MM state parameters recovered for the three benchmark datasets. For each generated dataset, the table lists the mpH$^2$MM states fitted using the ground-truth number of states, giving for each state its FRET efficiency (E), stoichiometry (S), and assignment (a FRET state or the donor-only state), together with the transition rates to every other state (e.g., k to S0, k to S3, in $s^{-1}$).
